# Disease mutation study identifies essential residues for phosphatidylserine flippase ATP11A

**DOI:** 10.1101/2020.01.13.904045

**Authors:** Kuanxiang Sun, Wanli Tian, Wenjing Liu, Yeming Yang, Xianjun Zhu

## Abstract

PS flippase (P4-ATPase) transports PS from the outer to the inner leaflet of the lipid bilayer in the membrane to maintain PS asymmetry, which is important for biological activity of the cell. ATP11A is expressed in multiple tissues and plays a role in myotube formation. However, detailed cellular function of ATP11A remains elusive. Mutation analysis revealed that I91, L308 and E897 residues in ATP8A2 are important for flippase activity. In order to investigate the roles of these corresponding amino acid residues in ATP11A, we assessed the expression and flippase activity of the respective ATP11A mutant proteins. ATP11A mainly localizes to the Golgi when co-expressed with TMEM30A, the β-subunit of the complex. Y300F and D913K mutations affect correct Golgi localization and PS stimulated flippase activity. In addition, Y300F mutation causes reduced ATP11A expression. Our data provides insight into essential residues for expression and flippase activity of ATP11A.

## 1. Introduction

Phospholipids of eukaryotic cell membranes are asymmetrically distributed, with phosphatidylserine (PS) and phosphatidylethanolamine (PE) concentrated mainly in the cytoplasmic leaflet of the membrane bilayer, while phosphatidylcholine (PC) and sphingolipids mainly positioned only on the exoplasmic leaflet [1–8]. Numerous physiological and biochemical processes, such as membrane stability and dynamics, cell polarity and migration, protein trafficking, cell apoptosis, cell signaling, platelet activation, neurodevelopment, blood coagulation and sperm cell capacitation, are relied on the asymmetric distribution of phospholipids [4, 9–19]. Mounting evidence reveals that the asymmetrically distributed phospholipids are maintained by P4-ATPases (also called flippases), which translocate phospholipids from the exoplasmic leaflet to the cytoplasmic leaflet [1, 5–8, 20]. In mammals, 14 members of P4-ATPases, designated ATP8A1 through ATP11C, have been identified [21]. ATP8A1, ATP8A2, ATP8B1, ATP8B2, ATP8B4, ATP10A, ATP10D, ATP11A, and ATP11C are localized to the plasma membrane, whereas ATP9A, ATP9B, ATP10B, and ATP11B are localized to intracellular membranes [1, 6, 9, 18, 22, 23]. Among the cell surface-localized P4-ATPases, ATP8A1, ATP8A2, ATP8B1, ATP11A, and ATP11C have been shown to catalyze the inward translocation of PS at the plasma membrane [6, 9, 18, 22, 23]. P4-ATPases, similar to other P-type ATPases such as Na^+^, K^+^-ATPase, are depending on heterodimeric interaction with a β-subunit CDC50 to proper folding and trafficking [11, 24–28]. In mammalian genome, there are three isoforms of CDC50 family: *CDC50A*, *CDC50B* and *CDC50C*, (also named *TMEM30A*, *TMEM30B* and *TMEM30C*, respectively). TMEM30A interacts with the vast majority of P4-ATPases, including ATP11A, and is critical for its proper functions [28, 29].

Thus far, the physiological functions of the majority of mammalian P4-ATPases were still unclear. Only mutations of several P4-ATPases were reported to cause severe human disease[30]. For instance, mutations identified in ATP8B1 are associated with liver disorders such as progressive familial intrahepatic cholestasis and hearing loss [31, 32]. Mutations in ATP8A2 cause severe neurological disorders such as cerebellar ataxia, mental retardation and disequilibrium syndrome [33–36]. ATP11C play a crucial role in differentiation of B cell in lymphopenia, anemia and intrahepatic cholestasis in mice [37–40]. ATP11A exhibits largely similar cellular distribution with ATP8A1 and ATP8A2[22, 41]. However, only a few studies were reported on ATP11A. ATP11A is ubiquitously expressed in various tissues [22] and deletion of *Atp11a* results in lethality during embryogenesis. Recent research indicated that the phospholipid flippase complex of ATP11A and CDC50A acts as a molecular switch for PIEZO1 activation that governs proper morphogenesis during myotube formation [42]. The detailed cellular function of ATP11A remains to be determined.

Previous study have reported *ATP8B1* mutations L127P and E981K in patients with familial intrahepatic cholestasis type 1 (PFIC1), while the mutation I344F is identified in patients with benign recurrent intrahepatic cholestasis type 1 (BRIC1) [43, 44]. An *ATP8B1* homozygous mutation L127V causes intrahepatic cholestasis in two Omani siblings [45]. The equivalent mutations of bovine ATP8A2 are I91P, I91V, L308F and E897K. To elucidate the functional consequences of flippase disease mutations, Rasmus H. et al. investigated the effect of mutations of those residues on expression and activity of ATP8A2 and found out the essential roles of these residues to the flippase translocation process [46]. Mutations of the I91 and L308 residues in ATP8A2 are positioned near proposed translocation routes in the protein. Mutation of the E897 residue is located at the exoplasmic loop between transmembrane helix 5 and 6. This mutational analysis suggested that I91, L308 and E897 residues affected the transport of phospholipids.

In order to investigate the effect of these above mentioned corresponding amino acid residues of ATP11A in the transporting phospholipids process, we matched these mutations to the equivalent sites of human ATP11A, which are I80P, I80V, Y300F and D913K respectively, and introduced those mutations into ATP11A ORF by site-direct mutagenesis technique. By investigating the expression pattern and flippase activity of these mutated ATP11A proteins, we demonstrated that variants of Y300F and D913K affected correct Golgi localization and the amount of PS internalization in plasma membrane. Additionally, Y300F mutation led to a decrease in ATP11A expression. This data provides insight into residues important for expression and flippase activity of ATP11A protein.

## Materials and methods

### Cell culture and transfections

HEK 293T and COS7 cells (American Type Culture Collection (ATCC, Manassas, VA, USA) were cultured in 4mM L-glutamine and 4500mg/ml glucose DMEM (HyClone) supplemented with 10% fetal calf serum (Gibco) and 1% antibiotics (HyClone) at 37℃ in 5% CO2 atmosphere. When HEK 293T cell density is 50% on 6 well plates and Cos7 cell density is 30%, 1 μg plasmid DNA was added with Lipofectamine 3000 (Invitrogen), 50 μl/ml OPTI-MEM (Gibco)/DMEM. Cells were harvested after 36-48 hours.

### Site-directed mutagenesis

Primers carrying the mutations were designed using NEBaseChanger software. Primer sequences: I80P-F: 5’-TTTCCTTATCCCATTTCTGGTGCAG-3’, I80P-R: 5’-TAAAAGTTGGCTACTCTTC-3’; I80V-F: 5’-TTTCCTTATCGTATTTCTGGTG-3’, I80V-R: 5’-TAAAAGTTGGCTACTCTTC-3’; Y300F-F: 5’-CTCATTGTGTTCCTCTGCATTCTG-3’, Y300F-R: 5’-GAACGCATTCATCGATTTTTC-3’; D913K-F: 5’-GACTTTGTACAAGACCGCGTATC-3’, D913K-R: 5’-TGTTGTGAAAACCCACAG-3’. Mutations were introduced into the ATP11A ORF using a The Q5® Sit-direct mutagenesis kit (New England Biolabs, Ipswich, MA, USA) following the manufacturer’s instruction. The mutated plasmids were sequenced to verify the successful introduction of the mutation.

### Immunoblotting

36-48 hours after transfection, HEK 293T cells were lysed in RIPA lysis buffer (150 mM NaCl, 50 mM Tris-HCl, 1%Triton X-100, 0.5% sodium deoxycholate, 0.1% SDS, pH 7.4) supplemented with Complete Protease Inhibitor Cocktail (Roche). The protein concentration of the lysates was determined using a DC Protein Assay (Bio-Rad). Equal amounts of protein (20ug) were loaded onto a 10% polyacrylamide gel and 0.45μM nitrocellulose membrane (Millipore, Billerica, MA, USA, catalog # HATF00010) was used for electrophoretic transfer. The blots were blocked with 8% non-fat dry milk in Tris-buffer saline with 0.1% Triton X-100 (TBST) for 2 hours at room temperature. Then, the membranes were incubated with the primary antibodies in blocking solution overnight at 4°C. The following primary antibodies were used for the Western blotting: mouse antibodies against Flag (Sigma-Aldrich, St. Louis, MO, USA) and mouse antibody against β-actin (Proteintech Group, Chicago, IL, USA). The primary antibodies were detected with anti-mouse HRP-conjugated secondary antibodies (1:5000; Bio-Rad, Hercules, CA, USA), and the signal was developed using Supersignal West Pico Chemiluminescent Substrate (Thermo Scientific). The relative intensity of the immunoreactive bands was quantified using the gel analysis tool provided in the ImageJ software. Normalization of the proteins of interest was performed relative to β-actin.

### Immunocytochemistry

36-48 hours after COS7 cells transfection, the transfected cells were washed twice with PBS, fixed with 4% paraformaldehyde and washed three times with PBS. Cells were then permeabilized and blocked with normal goat serum, Triton X-100, and NaN_3_ in PBS for 1 hour at room temperature. ATP11A protein was labeled with mouse anti-Flag antibody (1:2000, Sigma, Germany) and the endoplasmic reticulum (ER) was labeled with a rabbit anti-Calnexin antibody (1:1000, Cell Signaling Technology, CA, USA). The Golgi apparatus was labeled with a rabbit anti-GM130 antibody (1:1000, BD Biosciences, Mississauga, ON). The secondary antibodies used were Alexa 488 goat anti-rabbit IgG, Alexa 594 goat anti-mouse IgG (1:500, Invitrogen, USA). The images were captured under a Zeiss LSM 800 confocal scanning microscope.

### Flippase activity assay

36-48 hours after transfection, COS7 cells were washed thrice with PBS and treated with 0.25% trypsin. Cells were harvested by centrifugation and washed with HBSS (Gibco, USA). Cells were incubated with 10μM 1,2-dioleoyl-sn-glycero-3-phospho-L-serine-N-(7-nitro-2-1,3-benzoxadiazol-4-yl) (18:1 NBD-PS) (1:300, Sigma-Aldrich) for 15 minutes at 25°C. Subsequently, ice-cold HBSS supplemented with 2% bovine serum albumin (Sigma-Aldrich) was added to the cells for 10 minutes on ice to extract PS on the extracellular surface. This process was repeated 3 times. Subsequently, cells were centrifuged at 4 ℃ for 5 minutes at 300g and suspended in HBSS and analyzed with a flow cytometer (CytoFLEX, Beckman Coulter). Mean fluorescence intensity was calculated for each group.

### Statistical analysis

Statistical analysis was performed by one-way analysis of variance or student’s t-test using GraphPad Prism 6 software. The differences were considered statistically significant at P values <0.05. The quantitative data are presented as the mean ±SEM as indicated in the figure legends. All experiments were performed in triplicate and repeated at least twice.

## Results

### Conservative analysis of ATP11A mutational sites

Previous studies [43, 44] reported that the ATP8B1 mutational sites found in patients, L127, I344 and E897, are located in exons 4, 12, and 24, respectively. And these mutations are highly conserved among 10 different species (Fig. 1A, B). Comparative amino acid sequence alignment of other ATP11A proteins across different species revealed that the I80, Y300 and D913 mutations occurred also at highly conserved positions (Fig. 1A), which located in exons 3, 11, and 24, respectively (Fig. 1B). Topology of ATP11A has ten transmembrane helices and A, P and N domains and the C-terminal regulatory domain [47, 48] (Fig. 1C). Mutation sites of I80 and Y300 are located in the transmembrane region, while mutation sites of D913 is located at the exoplasmic loop between transmembrane helices 5 and 6 (Fig. 1C). These data demonstrated that the corresponding mutational sites in ATP11A are crucial for the conformational stability of its transmembrane structure.

**Figure 1.**
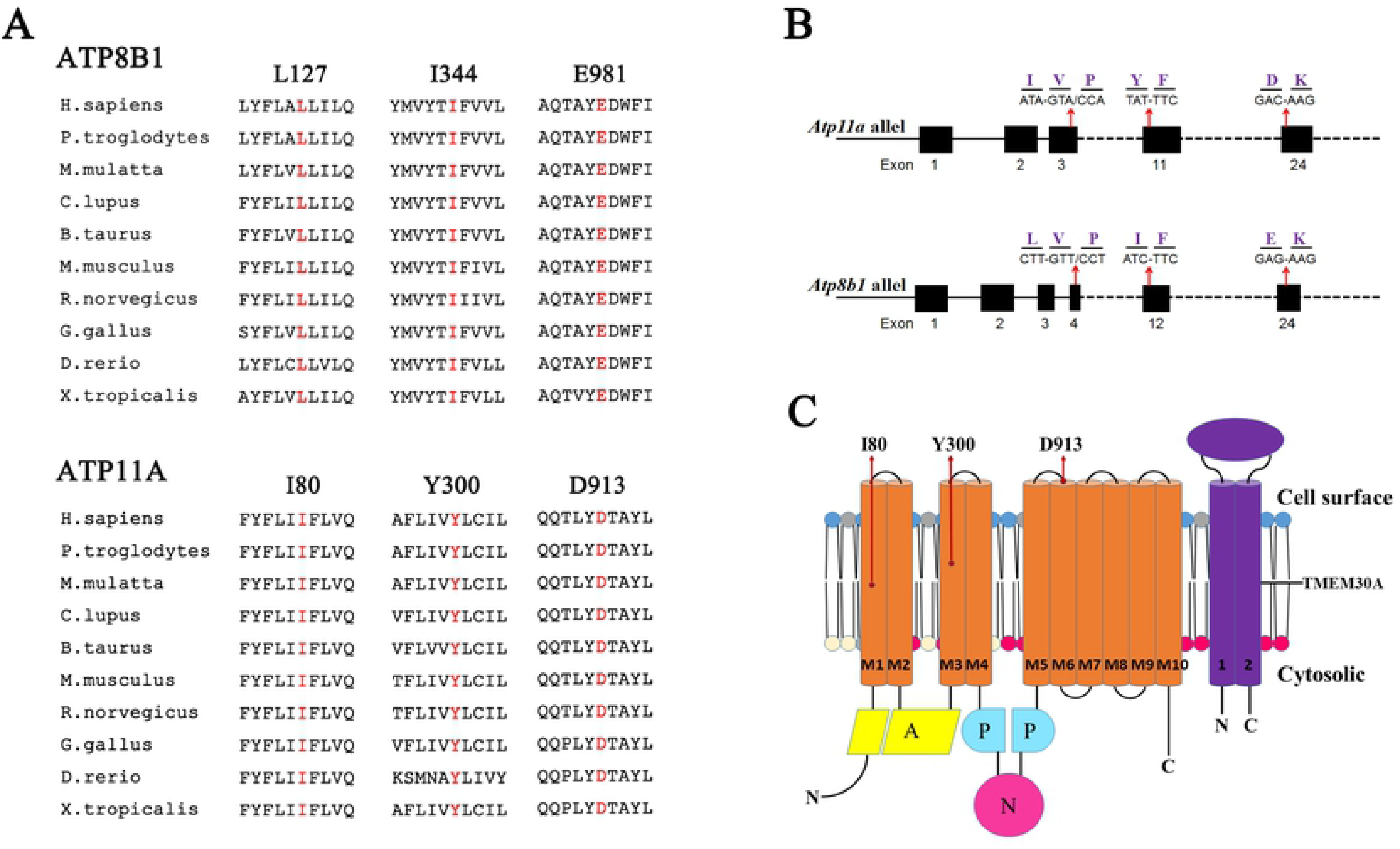
Location and conservation analysis of mutant residues. (A) ATP8B1 and ATP11A protein sequence alignment of amino acid sequences surrounding the identified mutations with its orthologues from *H. sapiens* to *X. tropicalis*. The mutated amino acid residues are in red. Note that the amino acid sequences of ATP11A surrounding the affected amino acid residues are highly conserved. (B) Schematic representation of exon-intron structure of *Atp8b1* and *Atp11a*. (C) Topology of ATP11A with the location of the mutations I80, Y300 and D913.

### The Y300F variant of ATP11A impairs its expression level

To investigate the expression levels of ATP11A variants, we transfected plasmids of ATP11A-WT, ATP11A-I80P, ATP11A-I80V, ATP11A-Y300F and ATP11A-D913K, which carried a C-terminal 3×Flag tag, into HEK 293T cells. Lysates from cells were separated by SDS-PAGE and subsequently subjected to immunoblotting analysis. The molecular weight of human ATP11A calculated from its primary structures is 129.7kDa (UniProtKB - P98196), whereas ATP11A protein was detected by immunoblotting with antibodies against Flag at approximately 145kDa (Fig. 2). Considering the membrane proteins generally undergo glycosylatation process before maturation [25, 49], the detected 145kDa ATP11A should be the mature glycosylated isoform. The immunoblotting data manifested that the expression level of ATP11A-Y300F in HEK 293T cells was reduced ~40% compared to ATP11A-WT (Fig. 2), while that of other ATP11A variants seems unchanged. The reduced content of ATP11A-Y300F likely resulted from the proteasomal protein degradation, which triggered by protein misfolding [50, 51].

**Figure 2.**
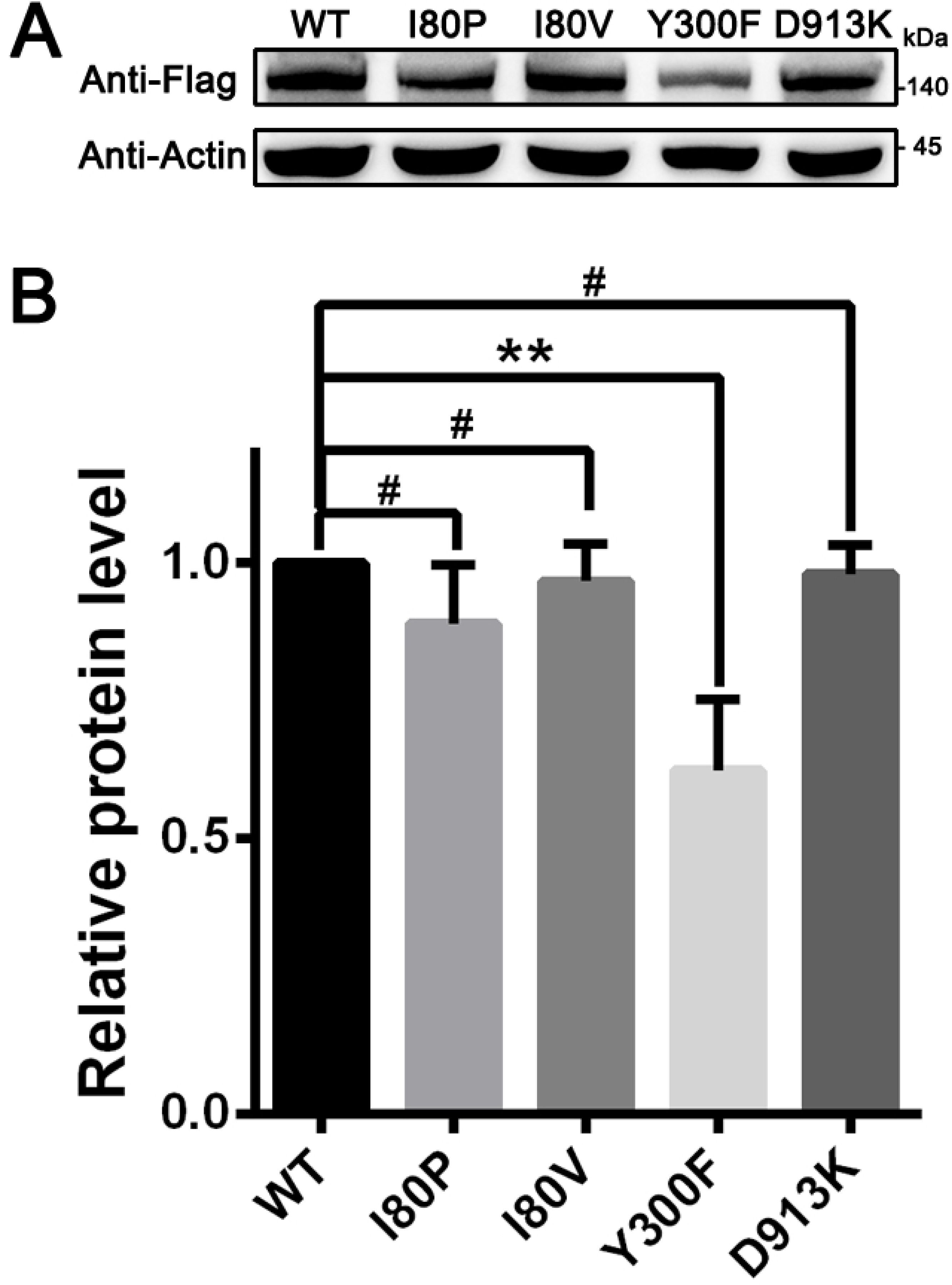
Expression of WT and mutant ATP11A proteins in HEK 293T cells. (A) Western blot analysis of ATP11A-WT, ATP11A-I80P, ATP11A-I80V, ATP11A-Y300F and ATP11A-D913K in HEK 293T cells using anti-Flag antibody. β-actin was used as the loading control. Uncropped gel pictures were shown in Figure S1. (B) Quantification of ATP11A revealed that expression level of ATP11A-Y300F was reduced. ATP11A-WT was used as control. N=3. **, p<0.01. The data represented means ±SEM. A representative result of three independent experiments was shown for A and B.

### The Y300F and D913K variants impairs the proper subcellular localization of ATP11A in the presence of TMEM30A

In order to detect the effect of mutations on the cellular localization of ATP11A, we first investigated the intracellular localization of variants by immunocytochemistry in the context of transfection of ATP11A without TMEM30A in COS7 cells. ATP11A-WT was largely localized to the endoplasmic reticulum (ER), which was manifested by the specific ER marker calnexin, and its mutants presented similar localization pattern (Fig. S1), suggesting little impact of these variants in ATP11A subcellular localization without the help of TMEM30A.

Previous studies have indicated that TMEM30A play a key role in the correct positioning of flippases [52, 53]. To this end, we further investigated the effect of TMEM30A co-expression on the subcellular localization of ATP11A by confocal microscopy. As mentioned above, when ATP11A and its mutants were single-transfected into COS7 cells, they were largely localized to ER (Fig. S1). However, in the presence of TMEM30A, large extent of ATP11A exit from ER to Golgi. This indicated that TMEM30A might help the correct positioning of ATP11A.

We further investigated the impact of variants on positioning with the help of TMEM30A. Notably, ATP11A-I80P and ATP11A-I80V mostly restricted to Golgi apparatus, while ATP11A-Y300F and ATP11A-D913K still resided at ER improperly (Fig. 3A). Statistical data manifested that the colocalization rate of ATP11A-WT, ATP11A-I80P and ATP11A-I80V with Golgi was about 60-70%, while that of ATP11A-Y300F and ATP11A-D913K with Golgi was only ~20-30% (Fig. 3B). These results demonstrated that ATP11A-I80P and ATP11A-I80V did not affect the correct sorting and targeting of mutant ATP11A proteins, whereas ATP11A-Y300F and ATP11A-D913K might obstruct the interaction with TMEM30A, resulting in the failure of positioning to Golgi apparatus with the help of TMEM30A.

**Figure 3.**
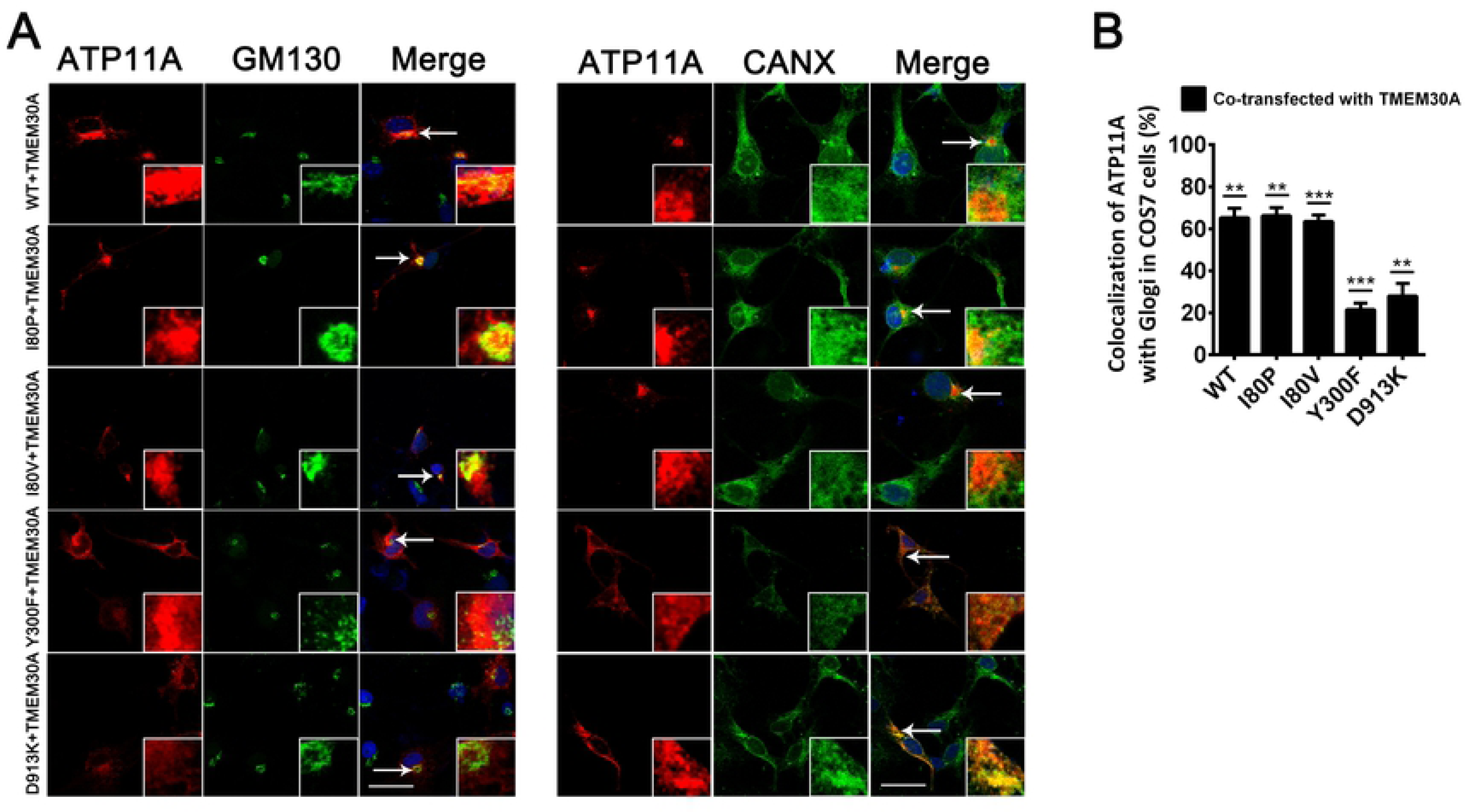
Subcellular localization of ATP11A mutants when co-transfected with TMEM30A in COS7 cells. (A) COS7 cells were transiently transfected with FLAG tagged ATP11A and HA-tagged TMEM30A plasmids. Cells were double stained for FLAG and organelle marker GM130 or Calnexin (CANX). Nuclei were counter-stained with DAPI (blue). Scale bars: 25 μm. (B) Percentage of colocalization of co-transfected ATP11A and TMEM30A with GM130-marked Golgi in COS7 cells. N=6. **, P<0.01; ***, P<0.001. Data are presented as the means ±SEM.

### The Y300F and D913K variants resulted in reduced PS internalization in plasma membrane

To investigate whether these mutations affect the phospholipid flippase activity, fluorescence-labeled PS analog (NBD-PS) was added into COS7 cells, which were transfected with the WT or variants plasmids of ATP11A. After incubation with NBD-PS for 15 minutes at room temperature, the cells were treated with 2% BSA to remove NBD-PS at the outer layer, and the fluorescence intensity of the inner layer was measured by flow cytometry (Fig. 4A-E). Compared to ATP11A-WT transfected cells, median NBD-PS fluorescence intensity (MFI) in ATP11A-Y300F and ATP11A-D913K transfected cells was decreased to 70% and 60% of the WT controls under the similar experimental conditions, suggesting reduced PS internalization in plasma membrane (Fig. 4F). These data demonstrated that variants of Y300F and D913K resulted in the reduced PS internalization in plasma membrane, which might be caused by a reduction in the amount of ATP11A or a decrease in the activity of ATP11A on the plasma membrane.

**Figure 4.**
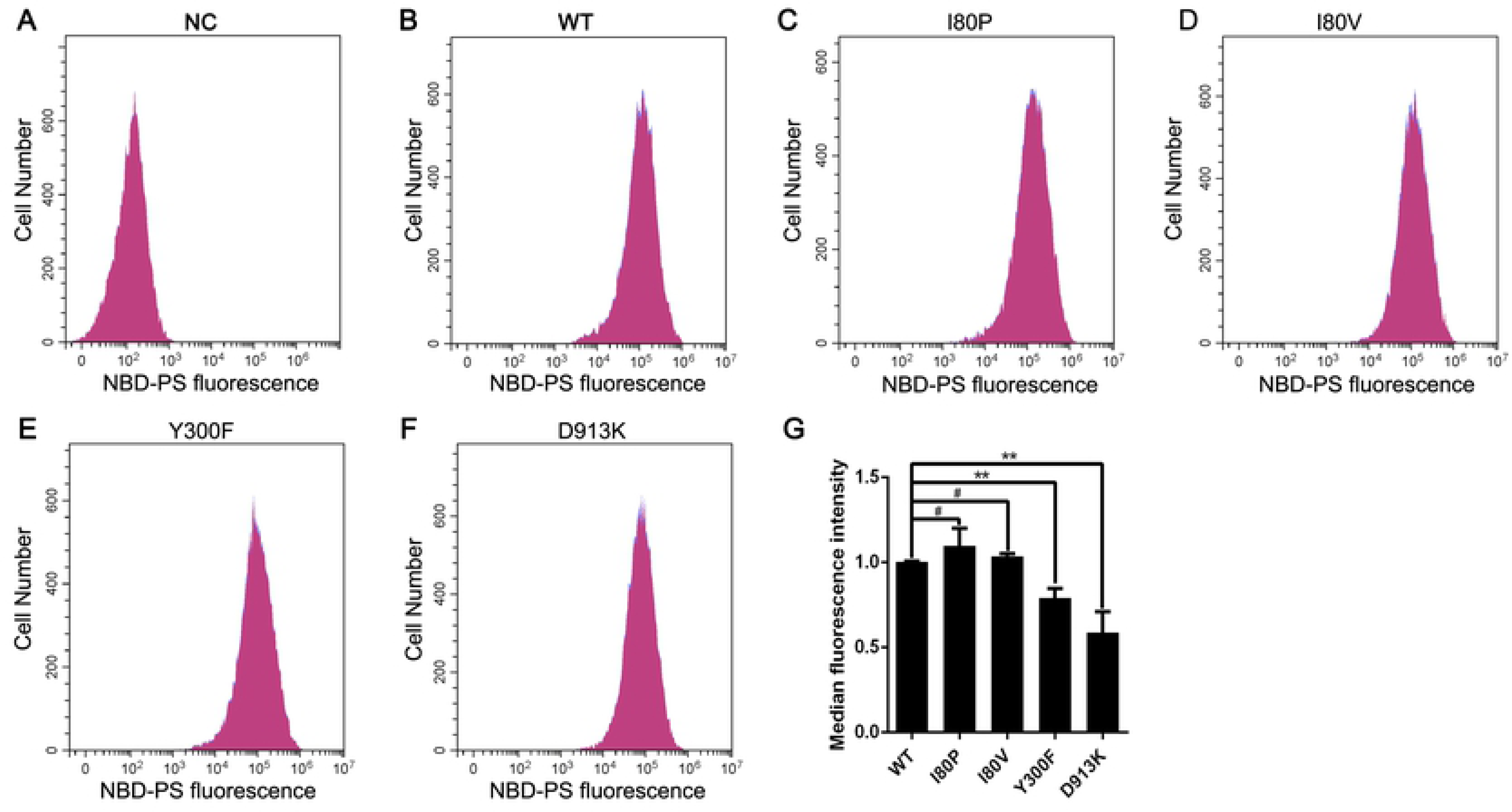
PS flippase activity assay of ATP11A mutants in COS7 cells. (A-G) NBD-PS labeling assays revealed partial loss of the amount of PS transported in ATP11A Y300F and D913K mutants. Flow cytometry histograms of NBD-PS labeling assays were shown in A-F. The X-axis represented NBD-PS fluorescence intensity of intact living cells. The Y-axis showed numbers of NBD-PS-labelled cells. (A) PcDNA3.1-C-3*flag was transfected as a negative control (NC). (F) PS flippase activity measured by mean fluorescence intensity of ATP11A mutants transfected cells. ATP11A-WT was used as control and its activity was set as 1.0. N=3. **, p<0.01. The data represented means ±SEM.

## Discussion

Eukaryotic P4-ATPases plays important roles in various cellular processes. Atp8a2 was reported to play a role in promoting neurite outgrowth in neuronal PC12 cells and rat hippocampal neurons [54]. Studies in mouse models have contributed to our understanding of the physiological functions of mammalian P4ATPases: *Atp8a1*-deficient mice exhibit delayed hippocampus dependent learning, *Atp8a2*-mutant mice display neurological abnormalities and *Atp11c*-deficient mice show arrested B-cell development [30]. However, little is known about the in vivo function of ATP11A. In order to get a deeper understanding of ATP11A, we selected three mutation sites L127, I344, E981 screened by ATP8B1, a member of the homologous family, as a reference. An ATP8B1 homozygous mutation ATP8B1-L127V causes intrahepatic cholestasis in two Omani siblings [45]. Mutation of ATP8B1-L127P has been reported to be related to PFIC1, but this mutant did not cause any change in the expression level and subcellular localization of ATP8B1 [43, 51]. ATP8B1-I344F detected in European families can cause BRIC1 [43]. A case of PFIC1 with mutation of ATP8B1-E981K in Japanese family was reported [44].

According to the amino acid sequence homology alignment, the equivalent positions of I80P, I80V, Y300F, D913K were determined in ATP11A. Mutant expression plasmids were constructed in vitro and its expression level, localization and activity of flippase in the cell were explored. We found that the expression of variant of Y300F was reduced by 40% by immunoblotting (Fig. 2), indicating that mutation at this site may lead to degradation of ATP11A protein. This mutation site is located in the third transmembrane domain (M3) of ATP11A protein (Fig. 1C) and is highly conserved in all 10 species sequences (Fig. 1A). In the crystal model, phenylalanine cannot be properly linked to isoleucine at position 359 on the M3-M4 loop (Fig. S2F). This indicates that mutations affecting this residue may lead to protein misfolding and eventually degradation by the proteasome.

In addition, the immunocytochemistry experiments demonstrated that variants of Y300F and D913K could not be correctly located in the Golgi apparatus when co-transfected with TMEM30A (Fig. 3A). Sites of Y300 and D913 are located at M3 and M5-M6 loop of ATP11A protein, respectively (Fig. 1C). In the crystal model, the variant Y300F causes the M3 and M3-M4 loop to be incorrectly connected (Fig. S2F), and variant D913K causes an error in the connection of M6 to M5-M6 loop (Fig. S2H). M3-M4 and M5-M6 loops are on the cell surface (Fig. 1C). It has been reported that in the extracellular region, the CDC50A ectodomain covers all of the extracellular loops of ATP8A1, except for the M1-2 loop, interacting in an electrostatic complementary manner: the extracellular loops of ATP8A1 bear negative charges, whereas CDC50A bears positive charges [47]. Therefore, we speculated that these two mutations interfere with the normal binding of ATP11A and TMEM30A and eventually cause ATP11A to be incorrectly located. Variants I80P and I80V only altered the connection of adjacent amino acids, but did not change the connection between the M1 and other loops (Fig. S2C, D). This also explains that these two variants have little effect on the ATP11A protein.

Flippase activity assay revealed that Y300F and D913K mutations resulted in reduced PS internalization in plasma membrane. These data demonstrated that variants of Y300F and D913K may resulted in the reduced PS internalization simply caused by reduced protein amounts in plasma membrane, because these mutant were not properly localized Golgi apparatus. These mutant also may directly reduce activity of the flippase in plasma membrane, eventually leading to a decrease in the amount of PS internalization [55, 56].

In summary, our data indicated that Y300F mutation of ATP11A could cause a decrease in ATP11A intracellular content, and variants of Y300F and D913K affected the subcellular localization and flippase activity of ATP11A. This indicates that the Y300 and D913 residues play an important role in the normal function of the ATP11A protein. Besides, we also verified that human TMEM30A protein is essential for the proper localization of ATP11A. These data provide basis for its pathogenic phenotype studies and give insight to its detailed cellular function.

## Acknowledgments

This study was supported by grants from the Department of Science and Technology of Sichuan Province (www.scst.gov.cn, 2016TD0009, 2017TJPT0010). The funders had no role in study design, data collection and analysis or preparation of the manuscript.

